# Non-retroviral endogenous viral element limits cognate virus replication in *Aedes aegypti* ovaries

**DOI:** 10.1101/2020.03.28.013441

**Authors:** Yasutsugu Suzuki, Artem Baidaliuk, Pascal Miesen, Lionel Frangeul, Anna B. Crist, Sarah H. Merkling, Albin Fontaine, Sebastian Lequime, Isabelle Moltini-Conclois, Hervé Blanc, Ronald P. van Rij, Louis Lambrechts, Maria-Carla Saleh

**Author notes:** These authors contributed equally (Yasutsugu Suzuki, Artem Baidaliuk). Unité de Parasitologie et Entomologie, Institut de Recherche Biomédicale des Armées (IRBA), Marseille, France; IRD, AP-HM, SSA, UMR Vecteurs – Infections Tropicales et Méditerranéennes (VITROME), IHU - Méditerranée Infection, Aix Marseille Université, Marseille, France. These authors contributed equally (Louis Lambrechts, Maria-Carla Saleh). Corresponding authors;, Address: 28 rue du Docteur Roux 75015 Paris, France.

## Abstract

Endogenous viral elements (EVEs) are viral sequences integrated in host genomes. A large number of non-retroviral EVEs was recently detected in *Aedes* mosquito genomes, leading to the hypothesis that mosquito EVEs may control exogenous infections by closely related viruses. Here, we experimentally investigated the role of an EVE naturally found in *Aedes aegypti* populations and derived from the widespread insect-specific virus, cell-fusing agent virus (CFAV). Using CRISPR/Cas9 genome editing, we created an *Ae. aegypti* line lacking the CFAV EVE. Absence of the EVE resulted in increased CFAV replication in ovaries, possibly modulating vertical transmission of the virus. Viral replication was controlled by targeting of viral RNA by EVE-derived piRNAs. Our results provide evidence that antiviral piRNAs are produced in the presence of a naturally occurring EVE and its cognate virus, demonstrating a functional link between non-retroviral EVEs and antiviral immunity in a natural insect-virus interaction.

## Introduction

Host genomes often harbor fragments of viral genomes, referred to as endogenous viral elements (EVEs), that are inherited as host alleles (Holmes, 2011). The best-studied EVEs are derived from mammalian retroviruses, which actively integrate their viral DNA into the host genome during their replication cycle. Retroviral EVEs play important roles in host physiology and antiviral immunity (Frank and Feschotte, 2017). Recent bioinformatic surveys also identified non-retroviral EVEs in a wide range of animal genomes, albeit their function was only studied in cell lines or protozoa (Belyi et al., 2010; Flynn and Moreau, 2019; Fujino et al., 2014; Horie et al., 2010; Katzourakis et al., 2014; Palatini et al., 2017; Parry and Asgari, 2019; Ter Horst et al., 2019; Waldron et al., 2018). The endogenization of non-retroviral sequences is presumably mediated by the activity of transposable elements (TEs), which are mobile DNA sequences ubiquitously found in eukaryotic genomes. Non-retroviral EVEs are often integrated in genomic regions surrounded by TEs, suggesting that TEs are involved in the integration and/or expansion of the EVEs (Gilbert and Feschotte, 2010; Horie et al., 2010; Palatini et al., 2017; Suzuki et al., 2017; Whitfield et al., 2017). The reverse transcription activity of retrotransposons is the likely mechanism generating non-retroviral DNA from RNA viruses, which are the hypothetical precursors of non-retroviral EVEs (Goic et al., 2016).

The recent discovery of non-retroviral EVEs in the genomes of mosquito vectors (Lequime et al., 2017; Palatini et al., 2017; Whitfield et al., 2017) has stimulated studies to elucidate their potential function. In particular, the genomes of the main arthropod-borne virus (arbovirus) vectors *Aedes aegypti* and *Aedes albopictus* harbor hundreds of non-retroviral EVEs predominantly derived from insect-specific viruses of the *Flaviviridae* and *Rhabdoviridae* families (Palatini et al., 2017; Whitfield et al., 2017). Interest in mosquito EVEs stems from the hypothesis that they may serve as the source of immunological memory against exogenous viruses in insects, as was recently reviewed in (Blair et al., 2020). This hypothesis largely relies on the observation that EVEs and their flanking genomic regions serve as templates for P-element-induced wimpy testis (PIWI)-interacting RNAs (piRNAs) (Palatini et al., 2017; Suzuki et al., 2017; Ter Horst et al., 2019; Whitfield et al., 2017). piRNAs are a major class of small RNAs (sRNAs) and are typically generated from genomic loci called piRNA clusters (Ozata et al., 2019). The piRNA pathway is considered a widely conserved TE-silencing system to prevent deleterious effects of transposition events in eukaryotic genomes, particularly in gonads (Cosby et al., 2019). In fact, production of EVE-derived piRNAs is observed across a wide range of animals such as mammals, arthropods and sea snails (Palatini et al., 2017; Parrish et al., 2015; Sun et al., 2017; Suzuki et al., 2017; Ter Horst et al., 2019; Waldron et al., 2018; Whitfield et al., 2017), in which EVEs are often enriched in piRNA clusters (Parrish et al., 2015; Russo et al., 2019; Ter Horst et al., 2019). The predominantly anti-sense orientation of EVE-derived piRNAs supports the idea that piRNAs could also mediate antiviral immunity by targeting exogenous viral RNA with high levels of sequence identity (Palatini et al., 2017; Russo et al., 2019; Whitfield et al., 2017).

The biogenesis of piRNAs and their function as a TE-silencing mechanism to protect genome integrity are well described in the model insect *Drosophila* (Czech and Hannon, 2016). piRNAs are characterized by their size of 26-30 nucleotides (nt) and distinctive sequence biases. Primary piRNAs typically display a uridine at the first nucleotide position, referred to as 1U bias. Secondary piRNAs overlap primary piRNAs over 10 nt at their 5’ extremity and display an adenine at their 10^th^ nt position, referred to as 10A bias (Brennecke et al., 2007; Gunawardane et al., 2007). These characteristics are a consequence of piRNA reciprocal amplification during the ping-pong cycle: (*i*) primary piRNAs are generated from single-stranded precursor RNA, (*ii*) primary piRNAs guide the cleavage of complementary RNA sequences, *(iii)* secondary piRNAs are generated from the 3’ cleavage products, *(iv)* secondary piRNAs induce cleavage of piRNA precursor transcripts, which are processed into primary piRNAs. Unlike *Drosophila,* it has been shown that mosquitoes produce virus-derived primary and secondary piRNAs during viral infections (Morazzani et al., 2012; Petit et al., 2016; Vodovar et al., 2012). Although most of these observations have been obtained using the *Ae. aegypti* cell line Aag2 and arboviruses such as dengue or Sindbis viruses, recent studies have shown that viral piRNAs were also found in mosquito cell lines persistently infected with insect-specific viruses, which are not infectious to vertebrates (Goertz et al., 2019; Rückert et al., 2019).

Whether EVEs can protect insects, and most importantly their germline, from viral infection through the piRNA pathway, has not been demonstrated *in vivo.* In mosquitoes, the antiviral activity of viral piRNAs is still debated and a direct link between EVEs and antiviral activity has yet to be established (Blair, 2019). One observation casting doubt on this hypothesis is that most arthropod EVEs identified so far are unlikely to serve as sources of antiviral piRNAs because they are not similar enough to currently circulating viruses (Parrish et al., 2015; Russo et al., 2019; Ter Horst et al., 2019). Here, we identified a new EVE in *Ae. aegypti* sharing ~96% nucleotide identity with a wild-type strain of cell-fusing agent virus (CFAV) that we previously isolated from *Ae. aegypti* in Thailand (Baidaliuk et al., 2019). CFAV is a widespread insect-specific virus infecting *Ae. aegypti* populations around the world (Baidaliuk et al., in press). We used this naturally occurring CFAV EVE and the cognate CFAV strain to experimentally investigate the antiviral function of mosquito EVEs in a natural insect-virus interaction. Analysis of sRNAs showed that the CFAV EVE produced primary piRNAs in the absence of CFAV infection. When mosquitoes were infected with CFAV, abundant CFAV-derived piRNAs were produced from the viral genomic regions overlapping with the CFAV EVE. piRNAs displayed a ping-pong signature as well as nucleotide biases consistent with production of EVE-derived primary piRNAs and virus-derived secondary piRNAs. Excision of the CFAV EVE by CRISPR/Cas9 genome engineering resulted in increased CFAV replication in ovaries. Our results provide empirical evidence that a non-retroviral EVE in *Ae. aegypti* contributes to the control of *in vivo* replication of a closely related exogenous virus via the piRNA pathway.

## Results

### Survey of CFAV-derived EVEs in *Aedes aegypti* genome sequences

In order to inventory CFAV-derived EVEs, we used BLAST search to identify CFAV-like sequences in publicly available *Ae. aegypti* genome assemblies, RNA sequencing data and whole-genome sequencing data. We identified several potential EVE structures based on samples for which reads aligned only to segments of the CFAV genome, in addition to samples for which reads covered the entire CFAV genome, presumably representing true CFAV infections (Figure S1). The predicted structure of two of these putative EVEs, which we designated CFAV-EVE1 and CFAV-EVE2, was obtained by *de novo* assembly (Figure 1A). These two putative EVEs were confirmed in an outbred *Ae. aegypti* colony derived from a wild population in Thailand and maintained in our laboratory since 2013. Using specific primer sets (Table S1), we detected CFAV-EVE1 and CFAV-EVE2 in 7 out of 8 and in 3 out of 8 individuals, respectively, in this outbred mosquito colony (Figure 1B).

**Figure 1.**
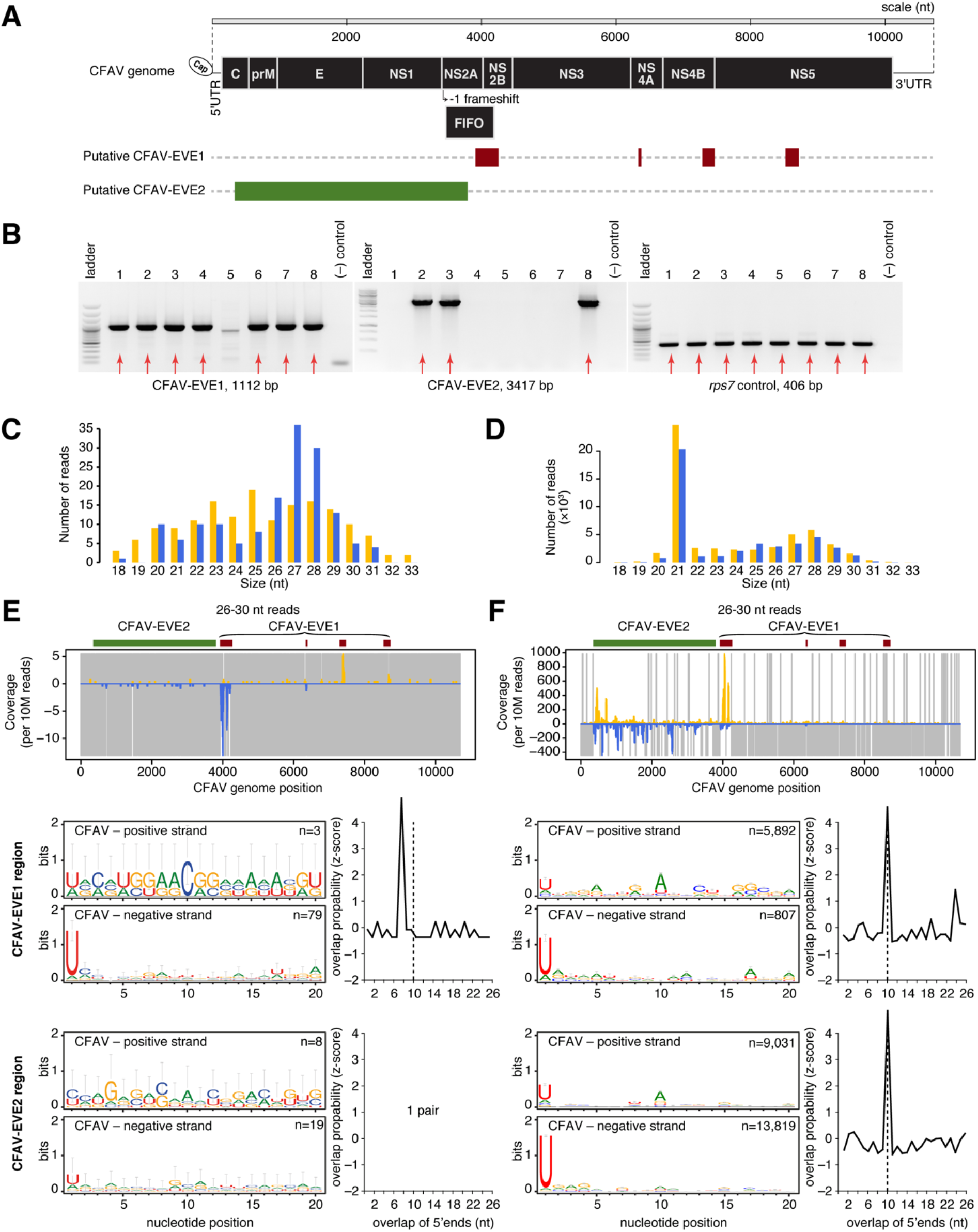
CFAV-derived endogenous viral elements interact with natural CFAV infection through the piRNA pathway. **A.** The schematic represents two potential CFAV EVE structures detected in publicly available *Ae. aegypti* sequences (See also Figure S1 and Table S2). **B.** The presence of putative CFAV-EVE1 and CFAV-EVE2 in eight mosquitoes from the same outbred colony was verified by PCR with primers specific to CFAV-EVE1 (left), CFAV-EVE2 (middle), and *rps7* gene control (right). **C-D.** Size distribution of sRNAs mapping to the CFAV genome from naturally CFAV-uninfected (**C**) and CFAV-infected (**D**) mosquitoes from the outbred colony. **E-F.** Analysis of CFAV-derived piRNAs from naturally CFAV-uninfected (**E**) and CFAV-infected (**F**) mosquitoes from the outbred colony. Mapping of 26-30 nt sRNAs (top), sequence logos of 26-30 nt sRNAs (bottom-left), and overlap probability of 26-30 nt sRNAs (bottom-right). Sequence logos and overlap probability for CFAV-EVE1 were restricted to the NS2 region. In panels **C-F**, positive- and negative-sense reads with respect to the reference CFAV genome are shown in yellow and blue, respectively. Uncovered nucleotides are represented by gray lines.

### CFAV-EVEs produce piRNAs that interact with viral RNA from a natural CFAV infection

Our outbred *Ae. aegypti* colony from Thailand is naturally infected with a wild-type strain of CFAV, which we previously isolated and named CFAV-KPP (Baidaliuk et al., 2019). Only a fraction of the mosquitoes in this colony are naturally infected, allowing us to investigate whether the CFAV EVEs produce piRNAs in the presence or absence of a natural CFAV infection. We sequenced sRNA libraries from both naturally infected and uninfected mosquito pools to examine sRNA production and specifically, EVE-derived and virus-derived piRNA production. In uninfected mosquitoes, the size distribution of the sRNA reads mapping to the CFAV-KPP genome sequence (Figure 1C) showed production of sRNAs of 26-30 nt in size with 1U bias, indicating that they are primary piRNAs generated mainly from the CFAV-EVE1 NS2 fragment, and to a lesser extent from the CFAV-EVE2 (Figure 1E). The lack of virus-derived 21-nt small interfering RNAs (siRNAs) confirmed the lack of CFAV infection in these mosquitoes (Figure 1C and Figure S2A). In contrast, the sRNA size profile of mosquitoes naturally infected with CFAV-KPP showed abundant production of virus-derived siRNAs (Figure 1D and Figure S2B). The CFAV-infected mosquitoes also harbored positive-stranded (+) CFAV-derived piRNAs, in addition to more abundant negative-stranded (-) primary piRNAs derived from both EVEs (Figure 1F) relative to the uninfected mosquitoes (Figure 1E). The presence of the 10A bias in (+) piRNAs and the 10-nt overlap probability between piRNA reads mapping to opposite strands was consistent with production of secondary virus-derived (+) piRNAs potentially triggered by EVE-derived (-) piRNAs, likely resulting in ping-pong amplification (Figure 1F). Thus, sRNA profiles in our outbred *Ae. aegypti* colony showed that the RNA transcribed from CFAV EVEs interacts with the viral RNA of a natural CFAV infection via the piRNA pathway.

### piRNAs from CFAV-EVE1 interact with viral RNA during CFAV experimental infection

To experimentally demonstrate the role of EVEs in antiviral immunity, we took advantage of a CFAV-free isofemale line of *Ae. aegypti* from Thailand maintained in our laboratory since 2010 (Fansiri et al., 2013; Lequime et al., 2016). We sequenced the whole genome of this isofemale line and only detected the presence of CFAV-EVE1 in the absence of other CFAV EVEs. CFAV-EVE1 was fully reconstructed from the newly obtained genomic data (Figure 2A, Table S2). The structure of CFAV-EVE1 in the isofemale line was consistent with the structure predicted from our bioinformatic survey (Figure 1A). CFAV-EVE1 consists of four adjacent fragments that correspond to the following CFAV genomic regions: NS5, NS4B-NS5, NS4A, and NS2A-NS2B/FIFO (designated as NS2 hereafter for simplicity). The CFAV-EVE1 sequence contains multiple start and stop codons in all six open reading frames. Moreover, two fragments (NS2 and NS4A) are inserted in opposite direction relative to the other EVE fragments, making it unlikely that functional viral peptides are effectively translated. We tested 31 individual mosquitoes from the isofemale line and found that 28 (90%; 95% confidence interval 73%-97%) were positive for CFAV-EVE1. As previously reported for other EVEs (Palatini et al., 2017; Suzuki et al., 2017; Ter Horst et al., 2019; Whitfield et al., 2017), CFAV-EVE1 and its flanking regions produced abundant antisense piRNAs (Figure 2B) when aligned to the isofemale line genome sequence. This observation indicates that CFAV-EVE1 is likely transcribed as a part of a longer piRNA precursor.

**Figure 2.**
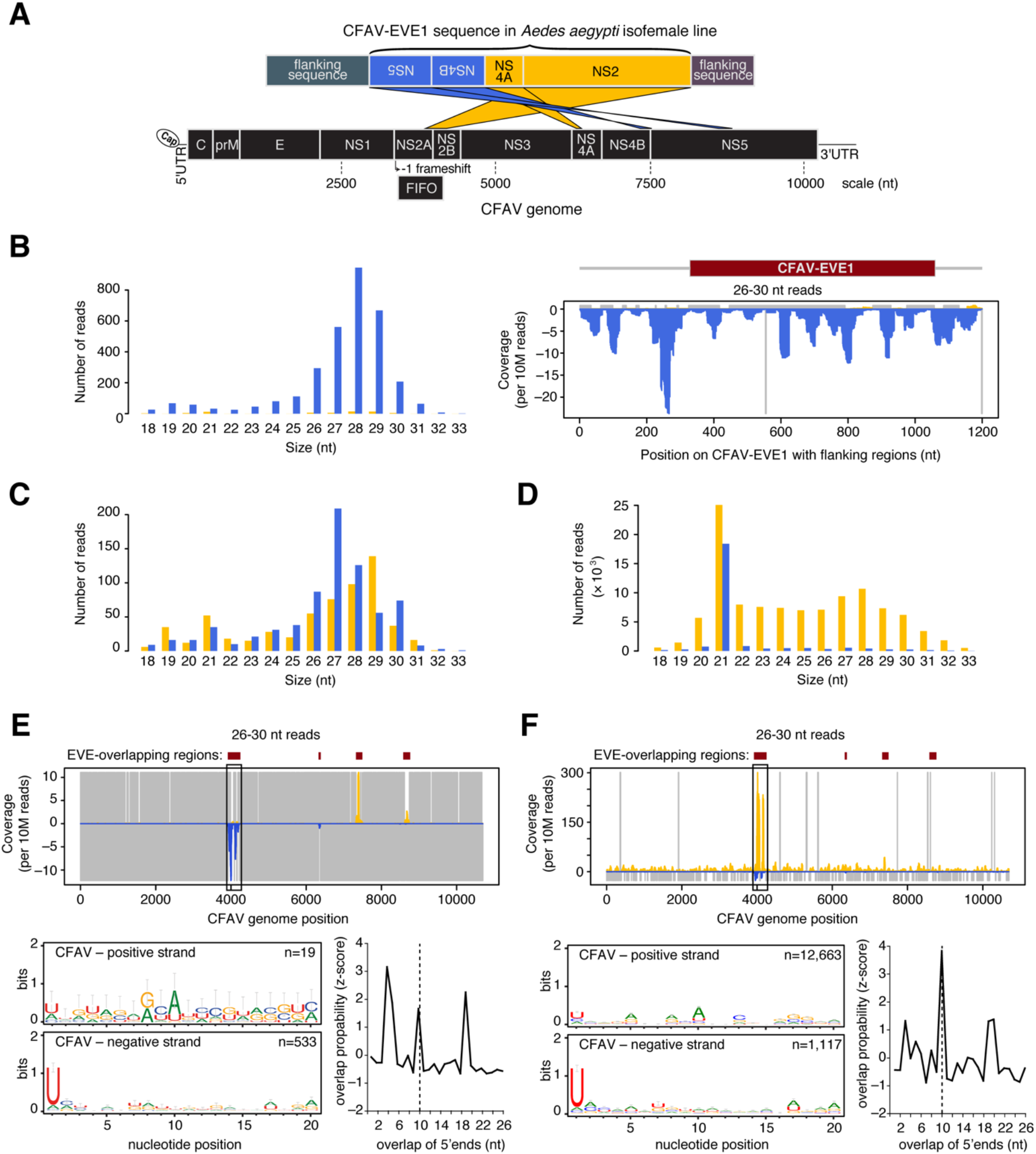
CFAV-EVE1 interacts with experimental CFAV infection through the piRNA pathway. **A.** Schematic of the CFAV-EVE1 structure in the CFAV-free isofemale line represented as the alignment of the EVE locus in the *Ae. aegypti* genome assembly AaegL3 (top) to the genome of the CFAV-KPP strain (bottom). CFAV-EVE1 comprises four different regions of the CFAV genome. Yellow and blue colors indicate forward and reverse strands, respectively, according to the transcription direction in the supercontig. **B.** Production of piRNAs from CFAV-EVE1 in the CFAV-free isofemale line, represented as the size distribution (left) and alignment to the CFAV-EVE-1 locus (right). Blue color corresponds to negative-sense reads with respect to the mapping reference. **C-D.** Size distribution of sRNAs mapping to the CFAV genome from experimentally CFAV-uninfected (**C**) and CFAV-infected (**D**) mosquitoes from the isofemale line. **E-F.** Analysis of CFAV-derived piRNAs from experimentally CFAV-uninfected (**E**) and CFAV-infected (**F**) mosquitoes from the isofemale line. Mapping of 26-30 nt sRNAs (top), sequence logos of 26-30 nt sRNAs (bottom-left), and overlap probability of 26-30 nt sRNAs (bottom-right). Sequence logos and overlap probability were restricted to the NS2 region. In panels **C-F**, positive- and negative-sense reads with respect to the reference CFAV genome are shown in yellow and blue, respectively. Uncovered nucleotides are represented by gray lines. See also Figure S2 and Figure S3.

The CFAV-EVE1 sequence of the isofemale line shared ~96% nucleotide identity with the CFAV-KPP genome, ranging from 94.6% to 98.8% among the different CFAV-EVE1 fragments (Table S2). To experimentally confirm our observations from naturally infected mosquitoes (Figure 1), we investigated the interaction between CFAV-EVE1 and the CFAV-KPP strain in the isofemale line (Figure 2C-D). In the absence of CFAV infection and as a consequence of the dual orientation of the CFAV-EVE1 fragments, EVE-derived piRNAs from the NS2 and NS4A regions were in antisense orientation, whereas EVE-derived piRNAs from the NS4B and NS5 regions were in sense orientation relative to the genome sequence of CFAV. We observed the most pronounced production of 1U biased, antisense primary piRNAs in the NS2 region (black frame in top panel of Figure 2E). When mosquitoes were inoculated with CFAV-KPP, the sRNA size profile (Figure 2D) showed abundant production of virus-derived siRNAs (21 nt) and also (+) CFAV-derived piRNAs corresponding to the CFAV-EVE1 genomic region of CFAV, in addition to (-) primary piRNAs derived from the EVE. As the NS2 region is the most abundantly covered by both sense and antisense piRNAs, we used this region (black frame in top panel of Figure 2F) to check for 10A bias as well as ping-pong signature. The 10-nt overlap of 5’ ends was consistent with active ping-pong amplification of the piRNAs in the NS2 region. In addition, analysis of the reads that unambiguously mapped to either the CFAV-KPP genome or to the CFAV-EVE1 sequence revealed that the vast majority of the piRNA reads derived from the CFAV-KPP genome were (+) piRNAs (Figure S3A) whereas almost all of the (-) piRNA reads derived from the CFAV-EVE1 (Figure S3B). It is worth noting that despite a similar abundance of EVE-derived primary piRNAs from the NS2 and NS4B regions in the absence of infection (Figure 2E), there is no evidence for amplification of piRNAs from the NS4B region during infection (Figure 2F). This suggests that the CFAV (-) RNA is not accessible or abundant enough for PIWI proteins loaded with primary piRNAs to initiate the ping-pong cycle.

Altogether, these results confirmed that CFAV-EVE1 produces piRNAs that target viral RNA and engage in a ping-pong cycle during experimental CFAV infection. The ability to experimentally infect the mosquito isofemale line carrying only CFAV-EVE1 with the CFAV-KPP strain allowed us to directly address the role of non-retroviral EVEs in antiviral immunity. This system recapitulated, under laboratory conditions, a unique situation found in nature (i.e., mosquitoes carrying an EVE that are infected or uninfected with a cognate virus).

### Genome engineering of a CFAV-EVE1 knockout line of *Aedes aegypti*

To directly test if the presence of CFAV-EVE1 influences CFAV replication in *Ae. aegypti,* we used CRISPR/Cas9 genome editing to create a CFAV-EVE1 knockout (−/−) line and a homozygous CFAV-EVE1 control (+/+) line derived from our CFAV-free isofemale line. We designed two single-guide RNAs (sgRNAs) targeting the boundaries of CFAV-EVE1 and another sgRNA in the middle of CFAV-EVE1 to promote excision (Figure 3A, Table S3). The sgRNAs were injected together with recombinant Cas9 into mosquito embryos. We obtained a heterozygous male devoid of CFAV-EVE1 (Figure 3B) that was outcrossed with wild-type mosquitoes from the parental isofemale line for two consecutive generations. The progeny were carefully sorted into purely CFAV-EVE1 homozygous (+/+) and knockout (−/−) mosquitoes. Importantly, the CFAV-EVE1 (−/−) mosquitoes only included the genetically engineered deletion genotype and excluded individuals that could be naturally devoid of CFAV-EVE1.

**Figure 3.**
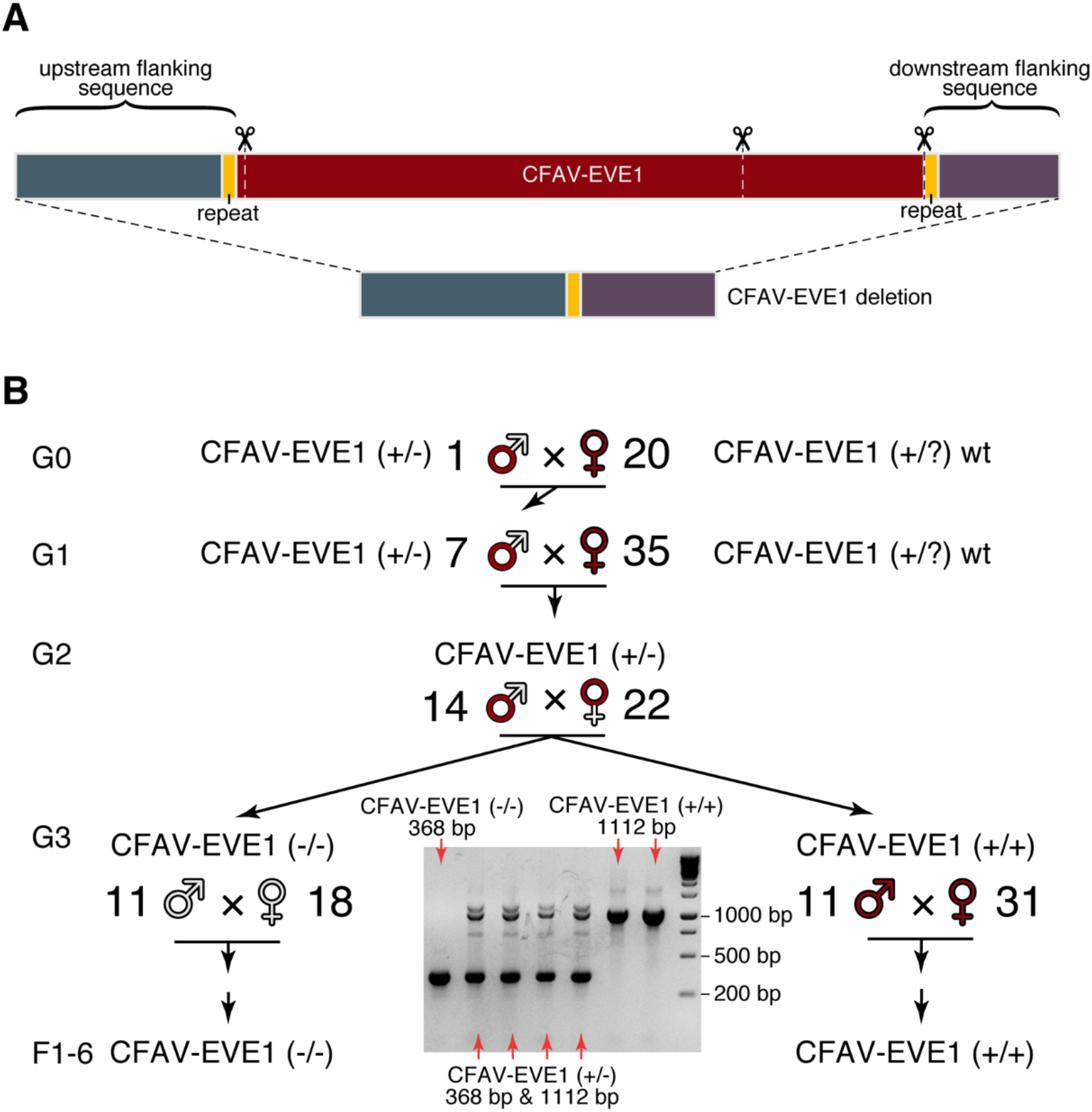
CRISPR/Cas9-mediated genome editing of CFAV-EVE1 in *Aedes aegypti*. **A.** Deletion of the CFAV-EVE1 from the *Ae. aegypti* genome of the CFAV-free isofemale line using CRISPR/Cas9. The upper bar represents the CFAV-EVE1 with the flanking regions and the three sgRNA target sites are shown with scissors. The lower bar represents the merged flanking regions without the CFAV-EVE1, where the short repeat sequences in the flanking regions (yellow segments on both bars) are merged into one. **B.** Generation of the CFAV-EVE1 (+/+) and (−/−) *Ae. aegypti* lines after CRISPR/Cas9-mediated genome editing. A single G0 male mosquito heterozygous for the CFAV-EVE1 deletion (+/−) was outcrossed with wildtype females harboring the CFAV-EVE1. The resulting heterozygous male G1 progeny was outcrossed with wild-type females harboring the CFAV-EVE1. The G2 heterozygotes of both sexes were intercrossed to produce a mixed G3 progeny that was sorted into pure homozygous CFAV-EVE1 (+/+) and (−/−) lines. The letter G denotes the generation of mosquitoes originating from the CFAV-EVE1 heterozygous male and wild-type females. The letter F denotes the generation of the CFAV-EVE1 homozygous lines. The agarose gel picture represents a fraction of samples genotyped at G3, where the pure homozygous individuals were selected by PCR genotyping of a single leg. See also Table S3.

### CFAV-derived piRNA production is strongly reduced in the absence of CFAV-EVE1

To determine if the absence of CFAV-EVE1 affected the production of CFAV-derived piRNAs, we infected CFAV-EVE1 (+/+) and CFAV-EVE1 (−/−) mosquitoes with CFAV-KPP. Seven days post infection, we dissected ovaries (germline tissue) and heads (somatic tissue) to prepare sRNA libraries from both tissues. Ovaries of mock-infected mosquitoes from the CFAV-EVE1 (+/+) line displayed the same sRNA profile (Figure S4A) as whole mosquitoes from the parental isofemale line (Figure 2C), with (-) piRNAs mainly derived from the NS2 region of CFAV-EVE1 and a 1U bias (Figure 2E and S4C). The heads of mock-infected mosquitoes (Figure S4E) contained few piRNAs mapping to the CFAV genome (<30 reads), consistent with the notion that germline tissues are the main producers of piRNAs (Akbari et al., 2013). As expected, mock-infected individuals from the CFAV-EVE1 (−/−) line did not harbor any piRNAs mapping to the CFAV genome in their ovaries and heads (Figure S4B, D, F, H). This result confirmed that genome editing effectively removed the CFAV-EVE1 sequence and allowed us to test whether the absence of the EVE affected the production of virus-derived piRNAs upon experimental CFAV-KPP infection. Of note, we detected a small number of viral siRNAs mapping to the CFAV genome in mock-infected heads of the CFAV-EVE1 (+/+) line (86 reads) and the CFAV-EVE1 (−/−) line (15 reads). As these samples were run in the same flow cell that contained CFAV-infected samples (Figure 4) producing thousands of viral siRNA reads in head tissues (31,988 reads in the CFAV-EVE1 (+/+) line and 8,465 reads in the CFAV-EVE1 (−/−) line), the minute amount of viral siRNA detected in mock conditions is likely due to demultiplexing cross contamination, a common and recurrent problem in high-throughput sequencing of multiplexed samples (Ballenghien et al., 2017).

**Figure 4.**
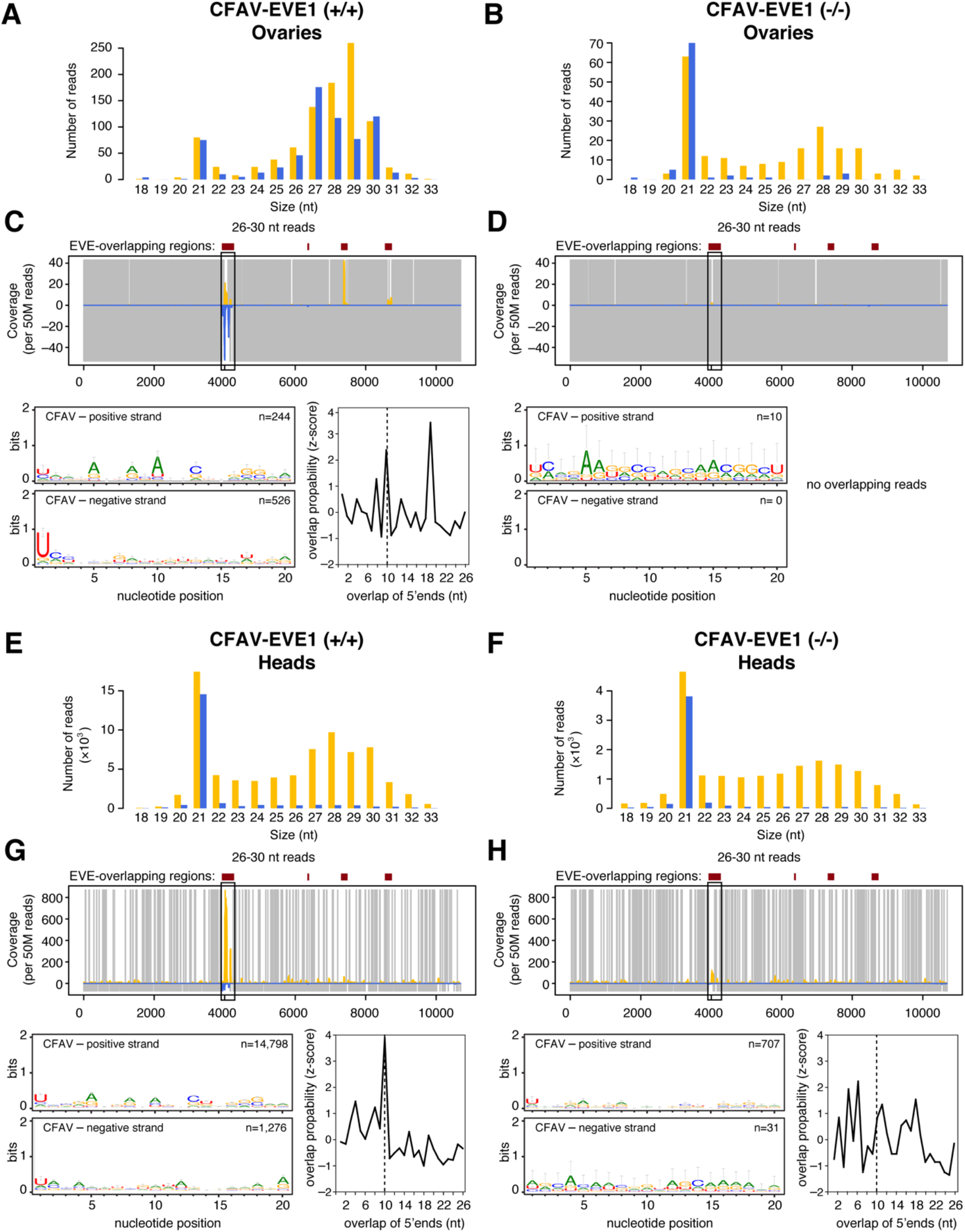
Ablation of CFAV-EVE1 prevents CFAV-derived piRNA amplification. Size distribution of sRNAs mapping to the CFAV genome in ovaries (**A-B**) and heads (**E-F**) from experimentally infected CFAV-EVE1 (+/+) (**A,E**) and CFAV-EVE1 (−/−) (**B,F**) mosquitoes 7 days post injection. Analysis of CFAV-derived piRNAs in ovaries (**C-D**) and heads (**G-H**) from experimentally infected CFAV-EVE1 (+/+) (**C,G**) and CFAV-EVE1 (−/−) (**D,H**) mosquitoes 7 days post injection. Mapping of 26-30 nt sRNAs (top), sequence logos of 26-30 nt sRNAs (bottom-left), and overlap probability of 26-30 nt sRNAs (bottom-right). Sequence logos and overlap probability were restricted to the NS2 region. In all panels, positive- and negative-sense reads with respect to the reference CFAV genome are shown in yellow and blue, respectively. Uncovered nucleotides are represented by gray lines. See also Figure S4 and Figure S5.

Following CFAV-KPP inoculation, we detected abundant viral siRNAs in both CFAV-EVE1 (+/+) and CFAV-EVE1 (−/−) mosquitoes (Figure 4A and Figure 4B). In addition, we detected virus-derived piRNAs and EVE-derived piRNAs with 1U and 10A bias and ping-pong amplification signature in the ovaries of CFAV-EVE1 (+/+) mosquitoes (Figure 4C). In contrast, (+) piRNAs mapping to the NS2 region of the CFAV-KPP genome were barely detectable in the ovaries of CFAV-EVE1 (−/−) mosquitoes (Figure 4D). Importantly, the detection of reads that unambiguously mapped to the virus showed that, even in the absence of the EVE, piRNAs were still produced from the virus genome upon infection (Figure S5A).

CFAV-KPP infection in the heads of CFAV-EVE1 (+/+) mosquitoes resulted in the production of CFAV-derived siRNAs as well as piRNAs (Figure 4E). The piRNAs corresponding to the NS2 region were in both sense and antisense orientation, presented a 1U-10A bias and 10-nt overlap of 5’ ends (Figure 4G). CFAV-KPP infection in the heads of CFAV-EVE1 (−/−) mosquitoes resulted in abundant CFAV-derived siRNAs (Figure 4F) and only piRNAs in sense orientation, without a ping-pong amplification signature, corresponding to primary piRNA production from the virus genome (Figure 4H and Figure S5B).

Altogether, these results showed that the production of CFAV-derived piRNAs is profoundly modified in the absence of CFAV-EVE1. Production of primary piRNAs from CFAV-EVE1 is necessary to trigger the production of secondary virus-derived piRNAs from the virus genome. This observation suggests that piRNAs could have an antiviral activity in the joint presence of an EVE and its cognate virus.

### Increased CFAV replication in ovaries in the absence of CFAV-EVE1

To assess the antiviral effect of piRNAs derived from the interaction between the EVE and the virus, we compared CFAV replication in CFAV-EVE1 (−/−) and CFAV-EVE1 (+/+) mosquitoes. To do so, we measured viral RNA levels in the heads and ovaries of females 4 and 7 days after CFAV inoculation. We performed six separate experiments using the same infectious dose and readout. The total amount of CFAV RNA produced by infected ovaries was significantly lower than the viral RNA produced in the heads (Figure 5). There was no consistent difference between mosquito lines across experiments for the CFAV RNA loads in heads collected on day 4 post inoculation (Figure 5A, Table S4). Accounting for the inter-experiment variation, there was a significant difference of CFAV RNA loads in ovaries on day 4 post inoculation, with CFAV replicating to higher levels in absence of the CFAV-EVE1 (Figure 5A, Table S4). On day 7 post inoculation, CFAV RNA loads were significantly higher in mosquito heads (Figure 5B, Table S4) and even more so in mosquito ovaries (Figure 5B, Table S4) in the absence of the CFAV-EVE1. Together, these experiments showed that CFAV replicated to higher levels in the absence of CFAV-EVE1, most prominently in ovaries. These results demonstrate the antiviral activity of an EVE against its cognate virus.

**Figure 5.**
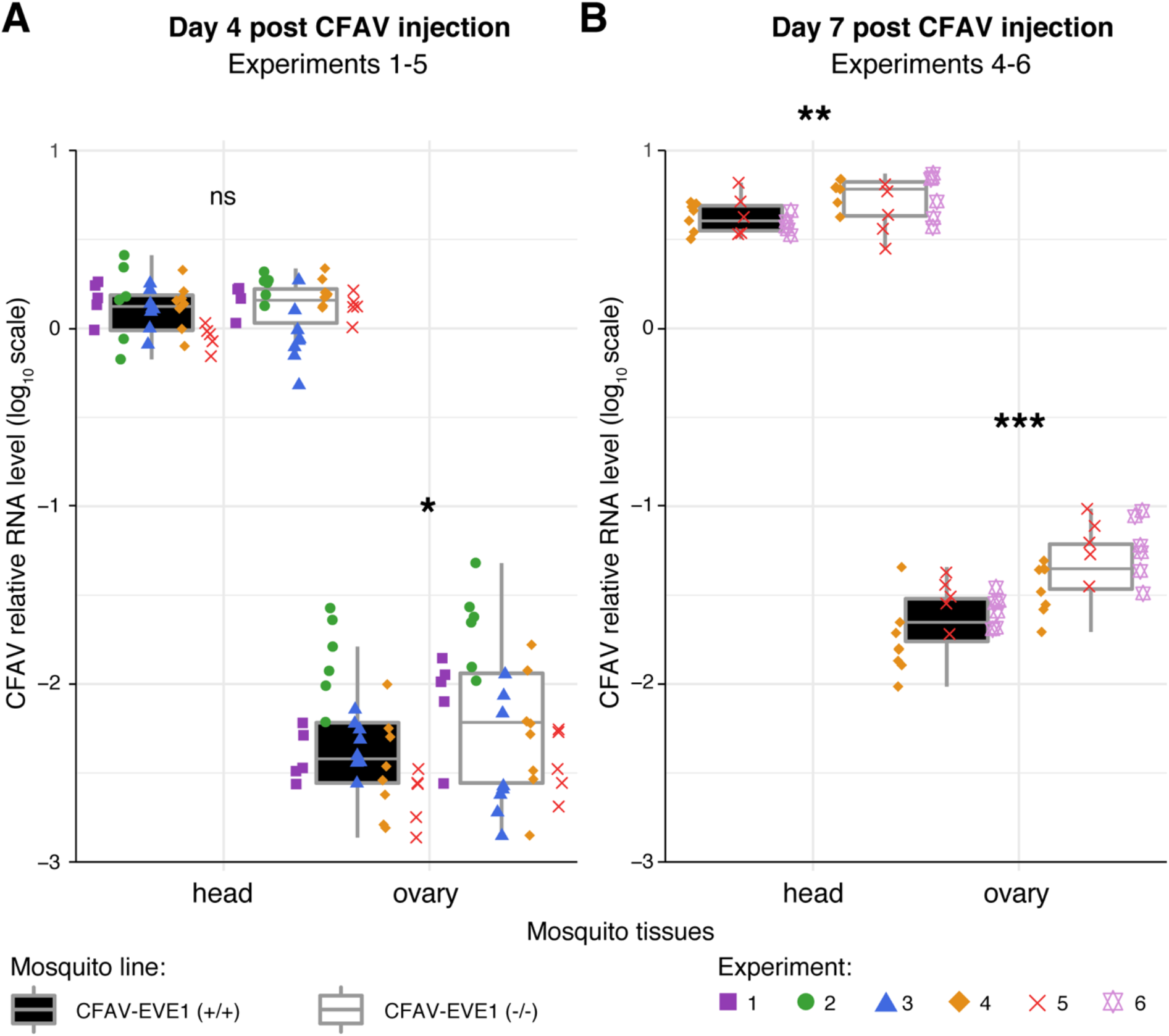
CFAV-EVE1 ablation results in increased CFAV RNA levels upon viral infection. **A-B.** Relative CFAV RNA levels (normalized by the *rp49* housekeeping gene) in heads and ovaries of the CFAV-EVE1 (+/+) (black boxplot) and CFAV-EVE1 (−/−) (white boxplot) *Ae. aegypti* lines on day 4 (**A**) and day 7 (**B**) post CFAV inoculation. Data are shown for six separate experiments represented by color- and symbol-coded data points. Relative viral RNA loads are represented by box plots in which the box denotes the median and interquartile range (IQR) and the whiskers extend to the highest and lowest outliers within 1.5 times the IQR from the upper and lower quartiles, respectively. Multivariate analysis of variance (MANOVA) was performed for each time point and tissue separately, accounting for the experiment, mosquito line and interaction effects. Stars indicate statistical significance of the mosquito line main effect accounting for the experiment effect (\**p* < 0.05, \*\**p* < 0.01, \*\*\**p* < 0.001, ns = not significant). The full MANOVA results are provided in Table S4.

## Discussion

The notion that non-retroviral EVEs could play a role in eukaryotic host immunity similar to retroviral EVEs (Anderson et al., 2000; Best et al., 1996) has recently gained traction. Several studies attempted to prove that non-retroviral EVEs contribute to the immune antiviral response. Perhaps the best example is Borna disease virus (BDV) and its endogenous bornavirus-like element, which affects BDV polymerase activity and inhibits virus replication in a mammalian cell line when incorporated into the viral ribonucleoprotein (Fujino et al., 2014). Tassetto *et al.* observed that mosquito cells carrying an EVE related to CFAV were partially protected against a recombinant Sindbis virus engineered to contain the EVE sequence (Tassetto et al., 2019). These *in vitro* experiments suggested that non-retroviral EVEs integrated in the host genome may provide antiviral protection against exogenous cognate viruses, but direct evidence from a natural system *in vivo* had not been provided until now.

The hypothesis that non-retroviral EVEs participate in antiviral immunity stems largely from accumulating evidence that they produce primary piRNAs (Palatini et al., 2017; Parrish et al., 2015; Sun et al., 2017; Suzuki et al., 2017; Ter Horst et al., 2019; Waldron et al., 2018; Whitfield et al., 2017). The piRNA pathway is often referred to as the guardian of genome integrity because its canonical function is to silence TEs in the germline (Czech et al., 2018). piRNA precursors are transcribed from genomic loci harboring transposon fragments that provide a genetic memory of past transposition invasion. The widespread occurrence of non-retroviral EVEs in *Aedes* mosquito genomes (Palatini et al., 2017; Whitfield et al., 2017) could reflect a similar mechanism whereby the function of EVEs would be to silence exogenous viruses with complementary sequences (Blair et al., 2020). A major challenge to prove this hypothesis is that the viruses currently circulating generally do not share a high nucleotide identity with the corresponding EVE sequences, preventing a possible match between EVE-derived piRNAs and the target viral RNA. In the present study, we overcame this obstacle by identifying a new EVE in *Ae. aegypti* mosquitoes from Thailand that is highly similar (~96% nucleotide identity) to a contemporaneous CFAV strain. We used this naturally occurring insect-virus interaction to test the hypothesis that a non-retroviral EVE can inhibit virus replication via the piRNA pathway *in vivo.*

Our results revealed that during both natural infection (mosquitoes carrying the CFAV EVE and naturally infected with CFAV) and controlled infection (mosquitoes carrying the CFAV EVE and experimentally inoculated with CFAV), the RNAs from the EVE and the virus interact through the piRNA pathway, resulting in inhibition of virus replication (Figure 6). Evidence of this interaction is provided by the abundant secondary piRNAs produced via the ping-pong amplification mechanism. Only when viral RNA is in presence of EVE-derived primary piRNAs does the piRNA pathway acquire its antiviral activity. Viral piRNAs alone are insufficient to induce this effect. Although viral piRNAs are commonly detected in mosquitoes (Miesen et al., 2016), their antiviral function has remained equivocal (Blair et al., 2020). Our study provides a clear demonstration that the piRNA pathway is involved in the mosquito antiviral response.

**Figure 6.**
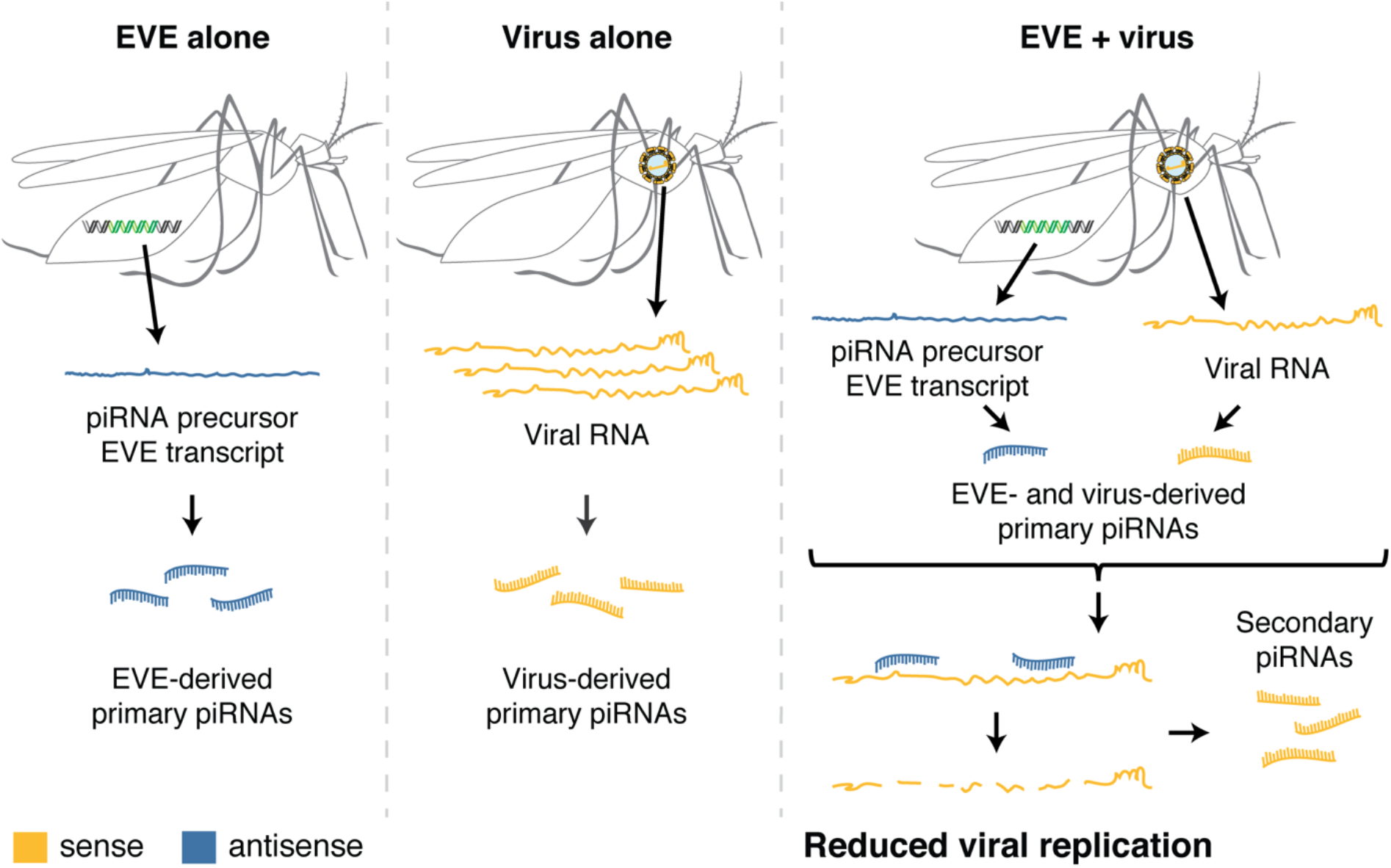
Model for the antiviral role of non-retroviral EVEs in mosquitoes. Both a naturally occurring EVE (left panel) and exogenous viral infection (middle panel) produce primary piRNAs, in antisense and sense orientation, respectively. Only when EVE and virus are present in the same mosquito, do piRNAs acquire antiviral activity (right panel) through EVE-derived piRNAs targeting the viral genome. Under this model, integration of non-retroviral sequences into the host genome, their transcription into piRNA precursors, and their processing into antiviral piRNAs are a mechanism by which EVEs confer heritable, sequencespecific host immunity.

We observed that the piRNA-mediated antiviral effect of the CFAV EVE was strongest in the ovaries. Although recent research on arthropods suggests that protecting the germline was not necessarily its ancestral role (Lewis et al., 2018), our results are consistent with a specialized role of non-retroviral EVE-mediated antiviral immunity in germ cells. Presently, little is known about the pathogenicity of insect-specific viruses in mosquitoes in nature. However, because they are thought to be primarily transmitted vertically from mother to offspring, it is likely that insect-specific viruses reduce fecundity and/or fertility of their host. We speculate that the EVE-piRNA pathway combination may have evolved to control the replication of vertically transmitted viruses in the germline and maintain high fecundity and fertility. In fact, minimizing the detrimental effects of viral infection in the germline benefits both the host and the virus because the fitness of vertically-transmitted viruses is directly linked to their host’s reproductive success (Anderson and May, 1982; Ewald, 1983, 1987).

Another open question is the degree of nucleotide identity required between the EVE and the virus for the antiviral activity to take place. Sequence mismatches reduce piRNA binding to its target sequences and it was shown that more than three mismatches can effectively abolish piRNA recognition of the target sequence in *Drosophila* (Huang et al., 2013). Even single mismatches in the seed sequence strongly reduces piRNA silencing efficiency in *Ae. aegypti* (Halbach et al., 2020). Therefore, viruses could escape EVE-mediated immunity by acquiring mutations, resulting in a possible coevolutionary arms race. Predicting the tempo and mode of such coevolutionary dynamics is difficult even when the fitness cost of individual mutations is known (Chabas et al., 2019). Interestingly, in our study the NS2 region of the CFAV EVE was most tightly involved in the interaction with the virus. This region corresponds to *fifo,* an open-reading frame (ORF) resulting from a ribosomal frameshift exclusively found in insect-specific flaviviruses (Firth et al., 2010). The existence of two overlapping ORFs in this region (main frame and −1 frame) thus constrains sequence evolution. We speculate that this region may have been specifically retained as an EVE in the *Ae. aegypti* genome because the high level of purifying selection in the *fifo* region may prevent CFAV from escaping the antiviral mechanism by sequence divergence.

In view of our results and the increasing body of evidence from the literature (Ophinni et al., 2019), we conclude that EVEs constitute a universal system of heritable, sequence-specific antiviral immunity in eukaryotes, analogous to CRISPR/Cas immunity in prokaryotes. In the particular case of mosquitoes, integration of non-retroviral sequences into the host genome, their transcription within piRNA clusters, and their processing into antiviral sRNAs constitutes a mechanism by which these acquired viral sequences are coopted to serve host immunity.

## Acknowledgements

We thank Catherine Lallemand for assistance with mosquito rearing. We are grateful to Catherine Bourgouin and Nicolas Puchot for assistance with the microinjection apparatus and to Anavaj Sakuntabhai for facilitation of the mosquito genome sequencing. R.v.R. was supported by the Netherlands Organization for Scientific Research (VICI grant 016.VICI.170.090). P.M. was supported by a short-term fellowship of the European Molecular Biology Organization (EMBO grant ASTF 449-2016). Work in the laboratory of L.L. was supported by Agence Nationale de la Recherche (grants ANR-16-CE35-0004-01 and ANR-17-ERC2-0016-01) and the City of Paris Emergence(s) program in Biomedical Research. Work in the laboratory of M.C.S. was supported by the European Research Council (FP7/2013-2019 ERC CoG 615220). L.L. and M.C.S were financed by the French Government’s Investissement d’Avenir program Laboratoire d’Excellence Integrative Biology of Emerging Infectious Diseases (grant ANR-10-LABX-62-IBEID). The funders had no role in study design, data collection and interpretation, or the decision to submit the work for publication.

## Author Contributions

A.B., Y.S., P.M., L.L., and M.C.S conceptualized the study. A.B. and Y.S. coordinated and performed infection experiments, analyzed and visualized the data, wrote the first draft and edited the manuscript. I.M.-C. participated in the infection experiments. H.B. performed sRNA sequencing. L.F. and P.M. analyzed and visualized sRNA sequencing data. A.B.C. generated genetically modified *Ae. aegypti* lines. S.H.M participated in the generation of the genetically modified mosquito lines. A.B. participated in the rearing of the genetically modified mosquito lines. A.F. conducted whole-genome sequencing and L.F. analyzed the whole-genome sequencing data. S.L. participated in the screening of the SRA database for CFAV-like sequences. R.v.R. participated in the interpretation of the results. L.L. and M.C.S supervised the study, provided resources, and edited the manuscript.

## Methods

### Ethics statement

Genetic modification of *Ae. aegypti* was performed under authorization number 4018 (bis) from the French Ministry of Higher Education, Research and Innovation.

### Survey of CFAV-related EVEs in public sequencing data of *Aedes aegypti*

The accession numbers for the *Ae. aegypti* sequencing data were selected using the web platform of the SRA database (Leinonen et al., 2011). We used BLAST (megablast) search (Altschul et al., 1990) implemented in the SRA Toolkit (Leinonen et al., 2011) to search for CFAV-like sequences in the preselected SRA data. The BLAST search resulted in 597 RNA-seq and 552 WGS runs tested, released before January 30 and February 6, 2020, respectively. Full-genome CFAV sequences from Thailand CFAV-Bangkok (European Nucleotide Archive, LR694074) (Zakrzewski et al., 2018) and CFAV-KPP (European Nucleotide Archive, LR596014) (Baidaliuk et al., 2019) were used as queries. Visualization of positive hits was performed in R v3.6.1 (http://www.r-project.org/). Using the online BLAST tool (megablast), the CFAV-EVE1 sequence was detected in the supercontig 1.109 of the AaegL3 genome assembly (GenBank accession number GCA_000004015.3) but absent from the AaegL5 genome assembly (GenBank accession number GCA_002204515.1). The CFAV-EVE1 sequence was reconstructed from a published WGS dataset (SRR5562867) using metaSPAdes v3.11.0 (Nurk et al., 2017). Reads from the WGS dataset were first quality trimmed with Trimmomatic v0.36 (Bolger et al., 2014) and aligned against the AaegL5 genome assembly with Bowtie2 v2.3.4.3 (--end-to-end --very-fast) (Langmead and Salzberg, 2012) to filter out all non-EVE reads. The CFAV-EVE2 sequence was reconstructed from SRA samples SAMN04480331, SAMN04480332,

SAMN04480333. Reads were trimmed with Cutadapt v1.18 (Martin, 2011). Relaxed local Bowtie2 v2.3.4.3 alignment (--local -D 20 -R 3 -L 11 -N 1 --gbar 1 --mp 3) was used in order to preselect CFAV-derived reads, which were then used for *de novo* assembly with Ray v2.3.1-mpi tool (Boisvert et al., 2010). The contigs obtained from all three SRA samples were combined into a single sequence of CFAV-EVE2 using Geneious (v10.2.3) software (https://www.geneious.com). The sequence was then verified by Bowtie2 alignment (--local) of the reads, coverage and single nucleotide variant calculation by bedtools v2.25.0 and LoFreq v2.1.3.1, respectively (Quinlan and Hall, 2010; Wilm et al., 2012). Both CFAV-EVE1 and CFAV-EVE2 sequences with annotations are available in Table S2.

### Live *Aedes aegypti* mosquitoes

#### Mosquito origin and maintenance

An outbred laboratory colony of *Ae. aegypti* mosquitoes originally sampled in 2013 from a wild population in Thep Na Korn Village, Kamphaeng Phet Province, Thailand (Lequime et al., 2016) was found to be infected with CFAV (Baidaliuk et al., 2019) and was used in this study for CFAV-EVE1 and CFAV-EVE2 detection by gDNA PCR and sRNA sequencing of naturally infected and uninfected mosquitoes. An isofemale line of *Ae. aegypti* originating from Kamphaeng Phet Province, Thailand was used for experimental infections *in vivo*. The isofemale line was created in 2010 as the progeny of a single-pair mating between a wild male from Mae Na Ree village and a wild female from Nhong Ping Kai village (Fansiri et al., 2013; Lequime et al., 2016). The inability to isolate CFAV from mosquito homogenates on C6/36 (*Ae. albopictus)* cells (ATCC CRL-1660) and negative RT-PCR directly on mosquito RNA confirmed that the isofemale line was CFAV-free. Mosquitoes were maintained under standard insectary conditions (27°C, 70% relative humidity and 12h:12h light:dark cycle). Larvae were reared in plastic trays filled with 1.5 L of dechlorinated tap water at a density of 200 larvae per tray and provided with 200 mg of TetraMin fish food (Tetra) on days 0 and 2 and 400 mg on day 4. After emergence, adult mosquitoes were housed in plastic cages under standard insectary conditions (27°C, 70% relative humidity and 12h:12h light:dark cycle) and provided with 10% sucrose solution *ad libitum.*

#### Whole-genome sequencing of the isofemale line

The whole genome of the *Ae. aegypti* isofemale line was sequenced at the 20^th^ generation of colonization. The DNA was extracted from a total of 144 virgin females following a published method (Bender et al., 1983). Six pools of 4 mosquitoes were homogenized in 240 μL of the following buffer: 0.1 M NaCl, 0.2 M sucrose, 0.1 M Tris buffer, 0.05 M EDTA, 0.5% SDS, pH adjusted to 9.2 with NaOH. The homogenates were incubated at 65°C for at least 35 min and 34 μL of 8 M KAc were added to the heated homogenates and cooled on ice for 30 min. Supernatants were transferred to new tubes, mixed with an equal volume of 100% ethanol and incubated for 5 min at room temperature (20-25°C). The DNA was pelleted by 15-min centrifugation at 21,100*g* and washed with 75% ethanol. The pellet was resuspended in 100 μL of PCR-grade water. This procedure was repeated 6 times and DNA elutes from all pools were gathered in a single tube and precipitated by adding 1/10 of 3M NaAc and 2.5x of cold 100% ethanol, followed by a washing step with 75% ethanol. The final elution was done in 400 μL of PCR-grade water. The genomic DNA was treated with RNase A/T1 (Thermo Scientific) for 30 min at 37°C and precipitated with NaAc again. The quality of the resulting DNA was assessed by Nanodrop (Thermo Scientific), Qubit HS DNA Assay Kit (Invitrogen), and 1% agarose gel migration. The DNA sequencing was performed commercially by Macrogen Europe (http://www.macrogen.com). A TruSeq PCR-free DNA shotgun library (550-bp inserts) was sequenced on an Illumina HiSeq 4000 platform (2 x 100 bp). The genome sequence of the isofemale line was deposited to Genbank (SRA sample SRR01437595).

#### DNA extraction and CFAV-EVE1-specific and CFAV-EVE2-specific PCRs

To verify the presence and prevalence of the CFAV-EVE1 in the *Ae. aegypti* isofemale line and outbred colony, DNA was extracted by two different methods. Genomic DNA was extracted from single legs of individual mosquitoes or whole individual mosquitoes using NucleoSpin DNA Insect Kit (Machery-Nagel) or NucleoSpin Tissue Kit (Macherey-Nagel) following the manufacturers’ instructions. Final elution was performed with 20 μL of the elution buffer. The DNA was used as a template for CFAV-EVE1-specific qualitative PCR with DreamTaq Green DNA Polymerase (Thermo Scientific) following the manufacturer’s recommendations, and using S7, EVE-GT-external, EVE-GTlong-external and/or EVE-GT-internal primers (Table S1). The CFAV-EVE2 sequence was detected with the CFAV-EVE2 primer set (Table S1).

Alternatively, DNAzol DIRECT (Molecular Research Center, Inc.) was used following manufacturer’s instructions, where DNA was extracted from single legs by placing a leg in 200 μL of DNAzol DIRECT in a 1.5-mL screw-cap tube partially filled with glass beads and homogenized. The lysate was centrifuged 15-30 sec at 21,100*g* and incubated at room temperature (20-25°C) for at least 20 min. Subsequently, 0.5-1 μL of lysate was used directly into a 20-μl PCR reaction. The same DNAzol DIRECT extraction procedure was used for whole mosquitoes, but the lysate was diluted 1:50 in PCR-grade water and 0.5-1 μL of the dilution was used in a 20-μL PCR reaction as described above.

### CRISPR/Cas9-mediated genome engineering

#### SgRNA design and synthesis

The *Ae. aegypti* isofemale line containing the CFAV-EVE1 was used to produce pure homozygous CFAV-EVE1 (+/+) and (−/−) lines using CRISPR/Cas9 as previously described for *Ae. aegypti* (Kistler et al., 2015). The single-guide RNAs (sgRNAs) were designed using CRISPOR (http://crispor.tefor.net/) by searching for 20-bp sgRNAs with the NGG protospacer-adjacent-motif (PAM). In order to reduce chances of off-target mutations, only sgRNAs with off-target sites which contained three or more mismatches were selected. Two sgRNAs with cut-sites proximal to the boundaries of the CFAV-EVE1 were chosen in order to delete the full CFAV-EVE1 sequence. A third sgRNA in the middle of the EVE sequence was added to facilitate deletion of the CFAV-EVE1 sequence. sgRNA sequences with their most probable off-target sites are represented in Table S3. SgRNAs were produced as previously described (Kistler et al., 2015). Double-stranded DNA templates for each sgRNA were produced by template-free PCR with two partially overlapping oligos (PAGE-purified, Sigma-Aldrich). Where necessary, one or two guanines were added to the 5’ end of the guide sequence within the primer to ensure the format “5’-GG(N18-20)-3’” in order to facilitate *in vitro* transcription with MEGAscript T7 *in vitro* transcription kit (Ambion). Transcribed sgRNAs were purified with MEGAclear kit (Invitrogen). Quality of sgRNAs were assessed with Bioanalyzer, Agilent 2100 Small RNA kit (Agilent).

#### Repair template design

We designed a 110-nt repair template with homology arms (HA) to the upstream and downstream flanking regions of the CFAV-EVE1 and extending to the sgRNA cut-sites (3 bp upstream of the PAM). The annotated sequence of the repair template is provided in Table S3. Due to the 5’ sgRNA having a cut-site inside the CFAV-EVE1 sequence, mismatches were artificially incorporated into to the 5’ HA of the repair template to ensure disruption of the CFAV-EVE1 sequence while maintaining enough homology to facilitate homologous recombination and deletion of CFAV-EVE1. An sgRNA sequence (with PAM) exogenous to the *Ae. aegypti* genome was also included in the repair template in an attempt to incorporate this guide sequence for further CRISPR/Cas9-mediated mutagenesis of this site. However, this and the modified CFAV-EVE1 sequences ultimately failed to get incorporated in CFAV-EVE1 (−/−) line genome. This could be explained by the presence of the 5’-TAAAAGTGGCGACGAG-3’ sequence contained in each flanking region of the CFAV-EVE1 that might have mediated the homology-dependent double-strand break repair independently of the repair template or that one homology arm acted as a truncated repair template.

#### Egg microinjection

The final microinjection mix contained 322 ng/μL spCas9 protein (New England Biolabs) with 40 ng/μL of each sgRNA and 127 ng/μL of the ssDNA repair template. The microinjection of *Ae. aegypti* embryos was performed according to standard protocols (Jasinskiene et al., 2007). *Ae*. *aegypti* females were engorged with commercial rabbit blood (BCL) via an artificial membrane feeding system (Hemotek). At least 3 days post blood meal, females were transferred into egg-laying vials and oviposition was induced by placing mosquitoes into dark conditions. Embryos were injected 30-60 min post oviposition. Embryos were hatched by being placed in water at least 3 days post injection and reared to adult stage as described above under mosquito maintenance. The generation 0 (G0) virgin adult mosquitoes were genotyped using a single leg DNA by PCR with EVE-GTlong-external primers (Table S1). The deletion in the CFAV-EVE1 heterozygous PCR products was confirmed by Sanger sequencing.

#### Generation of the CFAV-EVE1 (+/+) and (−/−) lines

A single male mosquito (G0) with a verified CFAV-EVE1 heterozygous genotype was mated with 20 wild-type females. The progeny (G1) were genotyped and 7 heterozygous males were mated with 35 wild-type females. G2 progeny were genotyped and 14 heterozygous males were mated with 22 heterozygous females. G3 progeny were genotyped and pure CFAV-EVE1 (+/+) and CFAV-EVE1 (−/−) lines were created by pooling homozygous positive (11 males and 31 females) and negative (11 males and 18 females) mosquitoes, respectively. The progeny of these crosses (F1) were verified by the PCR with CFAV-EVE1 external primers in 3 pools of 20 mosquitoes from each line. The lines were reared according to the standard rearing procedures described above. Further line genotype verification was performed at F3, F4, and F5. The F4 generation of mosquitoes was used for sRNA sequencing, which confirmed the almost complete absence of sRNAs complementary to CFAV in the CFAV-EVE1 (−/−) line, hence, the purity of the CFAV-EVE1 deletion and the absence of any other CFAV-related EVE that could have produced sRNAs.

### CFAV experimental infections *in vivo*

#### CFAV isolate and injection conditions

A wild-type CFAV strain (CFAV-KPP; ENA accession number LR596014) previously isolated from the *Ae. aegypti* outbred laboratory colony (Baidaliuk et al., 2019) was used for experimental infections of the CFAV-free *Ae. aegypti* isofemale line and the genetically modified lines. The first intrathoracic injection of the *Ae. aegypti* isofemale line harboring the CFAV-EVE1 was done with the third passage post isolation of the CFAV-KPP strain. The female mosquitoes were injected with 786 50% tissue-culture infectious dose (TCID50) units of virus per body using Nanoject II Auto-Nanoliter Injector (Drummond), then sacrificed on day 7 post injection and pooled RNA from 10 whole bodies was used for the first sRNA library preparation and sequencing. Mock injections were performed with naïve C6/36 cell-culture supernatant. The CFAV-KPP strain was also used for experimental infections of CFAV-EVE1 (+/+) and (−/−) lines (referred to as experiments 1-6), although it was produced from the viral genomic RNA instead of mosquito homogenates (Baidaliuk et al., 2019). Female mosquitoes were intrathoracically injected with 50 TCID50 units of CFAV-KPP per body in experiments 1-6 using Nanoject III Programmable Nanoliter Injector (Drummond). In experiment 5, mock injection was done with the naïve C6/36 cell-culture supernatant. RNA from the pools of heads and ovaries of injected females dissected on day 4 (experiments 1-5) or on day 7 (experiments 4-6) was used for the RT-qPCR with CFAV-specific primers and additionally for sRNA sequencing (experiment 5, day 7). In experiment 1, RNA was extracted from 5 pools of 4 tissues (pairs of ovaries or heads in all 6 experiments) per condition (mosquito line). In experiment 2, RNA was extracted from 6 pools of 5 tissues per condition (mosquito line). In experiment 3, RNA was extracted from 8 pools of 4 tissues per condition (mosquito line). In experiment 4, RNA was extracted from 6-8 pools of 4 tissues per condition (mosquito line and day post injection). In experiment 5, RNA was extracted from 5 pools of 9 tissues per condition (mosquito line and day post injection). Finally, in experiment 6, RNA was extracted from 5 pools of 5 tissues per condition (mosquito line). Mosquitoes were from generation F3 in experiments 1-4, generation F4 in experiment 5, and generation F6 in experiment 6.

#### CFAV RNA quantification

Total RNA was extracted and purified from mosquito tissues using TRIzol Reagent (Invitrogen) following manufacturer’s instructions with RNA elution in 30 μL of PCR-grade water. cDNA synthesis was performed using M-MLV reverse transcriptase (Invitrogen) by mixing 10 μL of eluted RNA with 100 ng of random primers (Roche), 10 nmol of each dNTP, 2 μL of DTT, 4 μL of 5X First-Strand Buffer, 0.5 μL of PCR-grade water, 20 units of RNaseOUT recombinant ribonuclease inhibitor (Invitrogen), and 200 units of M-MLV reverse transcriptase in a final reaction volume of 20 μL. Reactions were incubated for 10 min at 25°C, 50 min at 37°C, 15 min at 70°C, and held at 4°C until further use or stored at −20°C. cDNA was diluted 1:5 before quantitative analysis by qPCR was done using GoTaq qPCR Master Mix (Promega) following manufacturer’s recommendations. Primer sequences are provided in Table S2. CFAV qPCR values were normalized by the housekeeping gene *rp49* qPCR values and the normalized CFAV RNA levels were log_10_-transformed prior to their statistical analysis. A pool of mosquito tissues was considered a biological unit of replication. Type III multivariate analysis of variance (MANOVA) was performed separately for each time point (day 4 and day 7 post injection) and each tissue type (heads and ovaries). The linear model included experiment, mosquito line, and their interaction as covariates. The interaction term was removed from the model when its effect was statistically nonsignificant (*p* > 0.05), and type II MANOVA was performed instead. Statistical analyses were performed in the statistical environment R, version 3.5.2 (http://www.r-project.org/).

### Small-RNA sequencing

#### sRNA library preparation and sequencing

Total RNA from pools of 5 to 10 mosquitoes was subjected to acrylamide gel (15% acrylamide/bisacrylamie, 37.5:1, and 7M urea) electrophoresis to purify sRNAs of 19-33 nt in length. Purified sRNAs were used for library preparation with NEBNext Multiplex Small RNA Library Prep (Illumina) with 3’ adaptor, Universal miRNA Cloning Linker – S1315S (Biolabs) and in-house designed indexed primers. Libraries were diluted to 4 nM and sequenced on a NextSeq 500 sequencer (Illumina) with a NextSeq 500 High-Output Kit v2 (Illumina) (52 cycles).

#### Analyses of small-RNA sequencing data

The quality of the fastq files was assessed with FastQC software (www.bioinformatics.babraham.ac.uk/projects/fastqc/). Low-quality bases and adaptors were trimmed from each read using Cutadapt. Only reads with an acceptable quality (Phred score >20) and the adaptor sequence at the 5’ end were retained. A second set of graphics was generated by the FastQC software using the fastq files trimmed using Cutadapt. Reads were mapped to target sequences using Bowtie1 (one mismatch allowed between the read and its target for initial mapping or no mismatch allowed for target-specific mapping) or the Bowtie2 tool with default options for the sRNA or DNA library, respectively. The Bowtie1 tool (sRNA library) and the Bowtie2 tool (DNA library) generate results in sequence alignment/map (SAM) format. All SAM files were analyzed by the SAMtools package to produce bam indexed files. Homemade R scripts with Rsamtools and Shortreads in Bioconductor software were used for analysis of the bam files. For the analysis of sequence logos and sRNA overlaps, sRNA reads aligned to the CFAV-EVE1 sequence or to the CFAV genomic RNA were processed in Galaxy (Afgan et al., 2018). To generate sequence logos, reads of 26-30 nt in length were filtered and separated according to their genomic orientation. The selected reads were converted into FastA format, trimmed at the 3’ end to 20 nt and converted to RNA letters using the corresponding FastA/FastQ tools. The processed reads were used as input for the Weblogo tool available in the Galaxy toolshed (Crooks et al., 2004). For the analysis of ping-pong signatures, aligned reads were loaded into the Mississippi instance of Galaxy (https://mississippi.snv.jussieu.fr/). SAM files containing the reads of 26-30 nt in length were used as input for the Small RNA signatures tool. The Z-scores of the calculated overlap probabilities were plotted with Graphpad Prism. All sRNA sequencing library sizes with the number of CFAV-mapped reads are reported in Table S5. All data are available in the Sequence Read Archive repository under project PRJNA588447.

---

**Table S1. Primers used in this study.**

**Table S2. CFAV-EVE1 and CFAV-EVE2 sequence annotation based on similarity to the CFAV-KPP genome.** The CFAV-EVE1 sequence was reconstructed from the WGS data of the CFAV-free isofemale line from Thailand generated in this study. CFAV-EVE1 sequences extracted from the AaegL3 assembly and reconstructed from a published WGS dataset (SRA sample SRR5562867) are also provided. The CFAV-EVE2 sequence was reconstructed from published RNA-seq data of the lower reproductive tract of *Ae. aegypti* derived from Thailand (SRA samples SAMN04480331, SAMN04480332, and SAMN04480333).

**Table S3. CRISPR/Cas9 design for CFAV-EVE1 knockout.**

**Table S4. Analysis of variance of CFAV RNA levels in tissues of CFAV-infected CFAV-EVE1 (+/+) and (−/−) *Aedes aegypti* mosquito lines.** Multivariate analysis of variance (MANOVA) of relative CFAV RNA levels (normalized to the *rp49* housekeeping gene) was performed for each time point and tissue separately. The model included the effects of the experiment, the mosquito line and their interaction (when significant). The stars indicate the statistical significance of the effect (\**p* < 0.05, \*\**p* < 0.01, \*\*\**p <* 0.001). *Df* = degrees of freedom; *F* = *F* statistic; *p* = *p* value.

**Table S5. sRNA library information.**

