## Supplemental Figures for "Non-retroviral endogenous viral element limits cognate virus replication in *Aedes aegypti* ovaries"

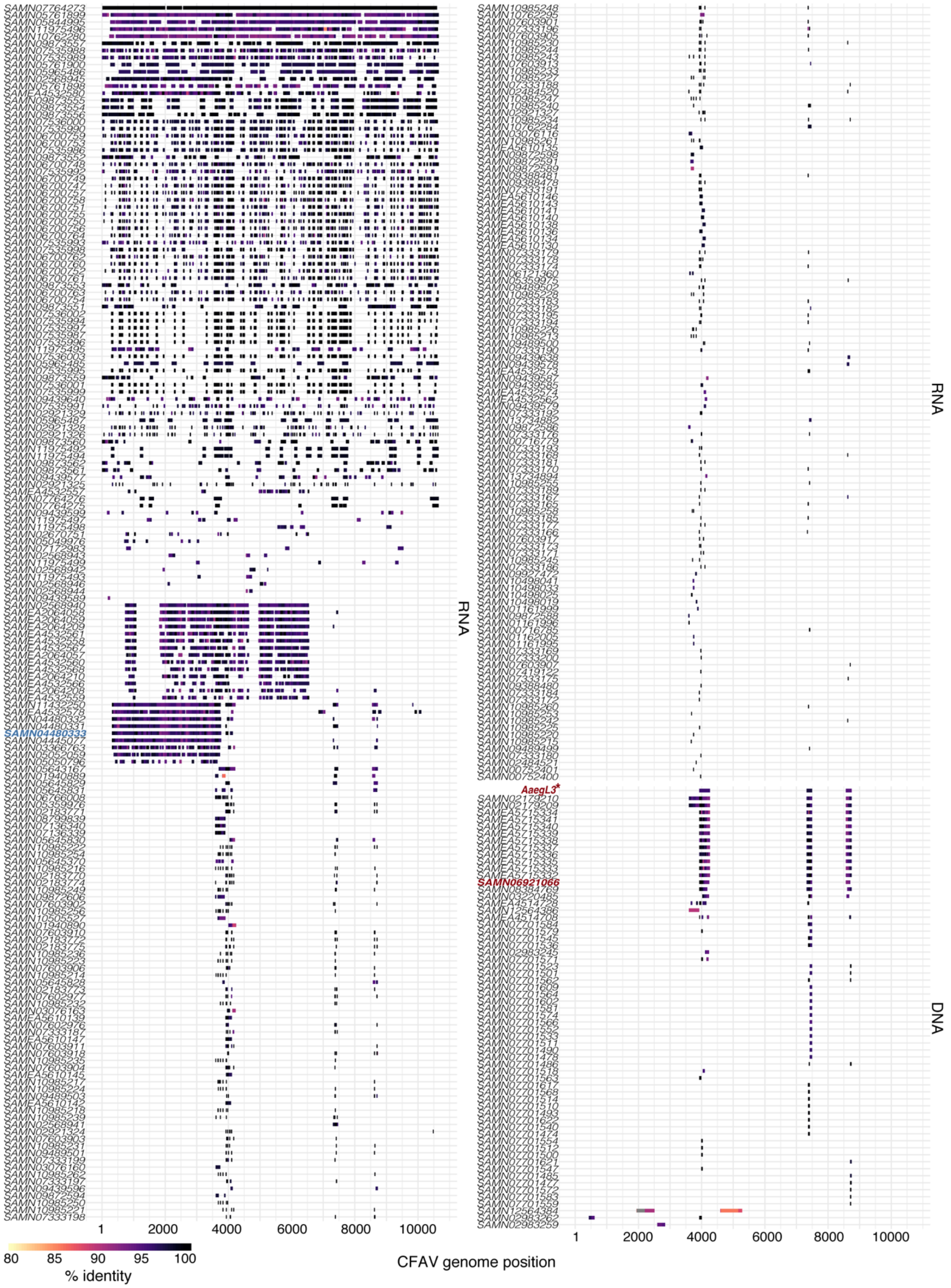


**Figure S1. BLAST alignment of publicly available *Aedes aegypti* sequencing data to the CFAV genome. Related to Figure 1A.**

Each line represents a BLAST hit from a Sequence Read Archive (SRA) RNA-seq sample, an SRA whole-genome sequencing (WGS) sample, or the AeagL3 assembly (indicated by an asterisk). BLAST hits are aligned to the CFAV genome and ordered according to the pattern of coverage. The percentage of nucleotide identity to the CFAV-Bangkok (RNA) or CFAV-KPP (DNA) genomes is indicated by the color gradient shown at the bottom of the plot. The sample names colored in red and blue were used to reconstruct the sequence of CFAV-EVE1 and CFAV-EVE2, respectively.

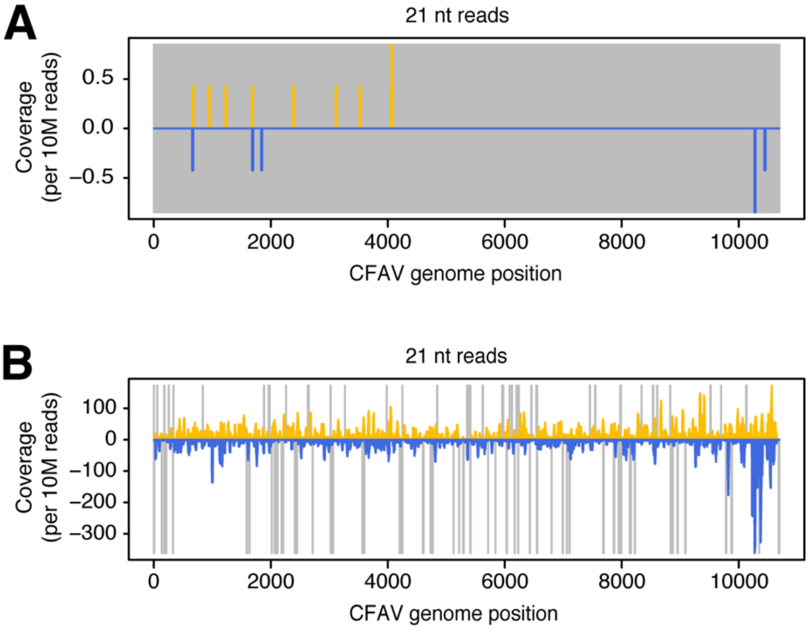


**Figure S2. siRNA response to a natural CFAV infection in *Aedes aegypti* mosquitoes harboring CFAV-EVE1 and CFAV-EVE2. Related to Figure 1.**

Profiles of siRNAs mapping to the CFAV-KPP genome from naturally CFAV-uninfected (**A**) or CFAV-infected (**B**) mosquitoes from the outbred colony. Positive- and negative-sense reads are shown in yellow and blue, respectively. Uncovered nucleotides are represented by gray lines.

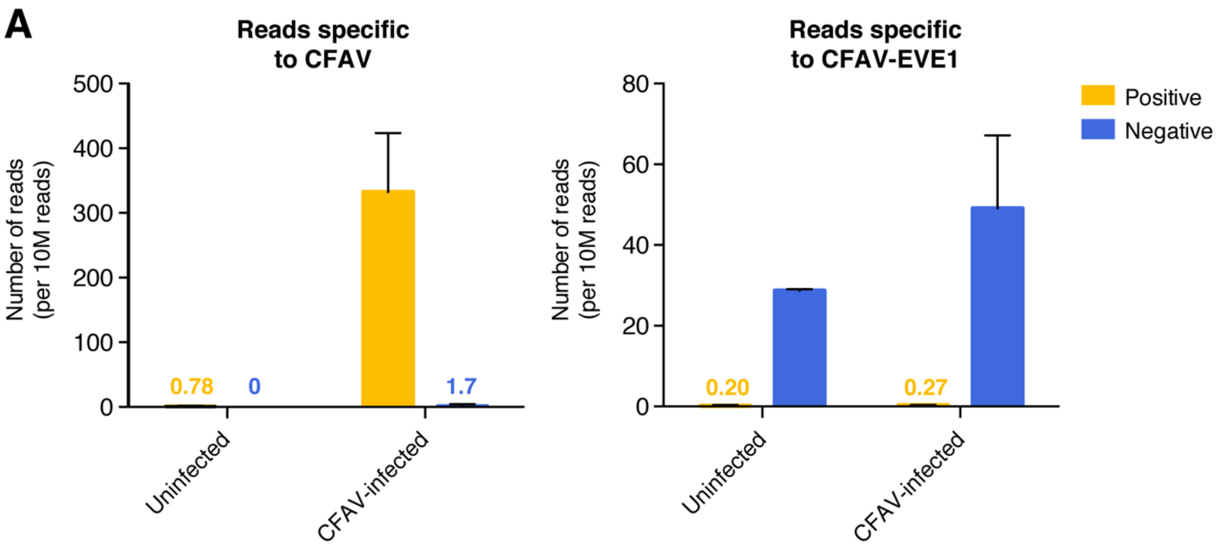


**Figure S3. Target specificity of piRNAs upon CFAV infection in the *Aedes aegypti* isofemale line harboring CFAV-EVE1. Related to Figure 2.**

Number of 26-30 nt sRNA reads unambiguously mapping to the CFAV-KPP genome (**A**) or to the CFAV-EVE1 locus (**B**) in uninfected or CFAV-infected individuals of the isofemale line. Yellow and blue colors represent positive-sense and negative-sense reads, respectively. For visual clarity, the normalized number of reads is shown above the bars when the number of reads is <1% of the maximum value.

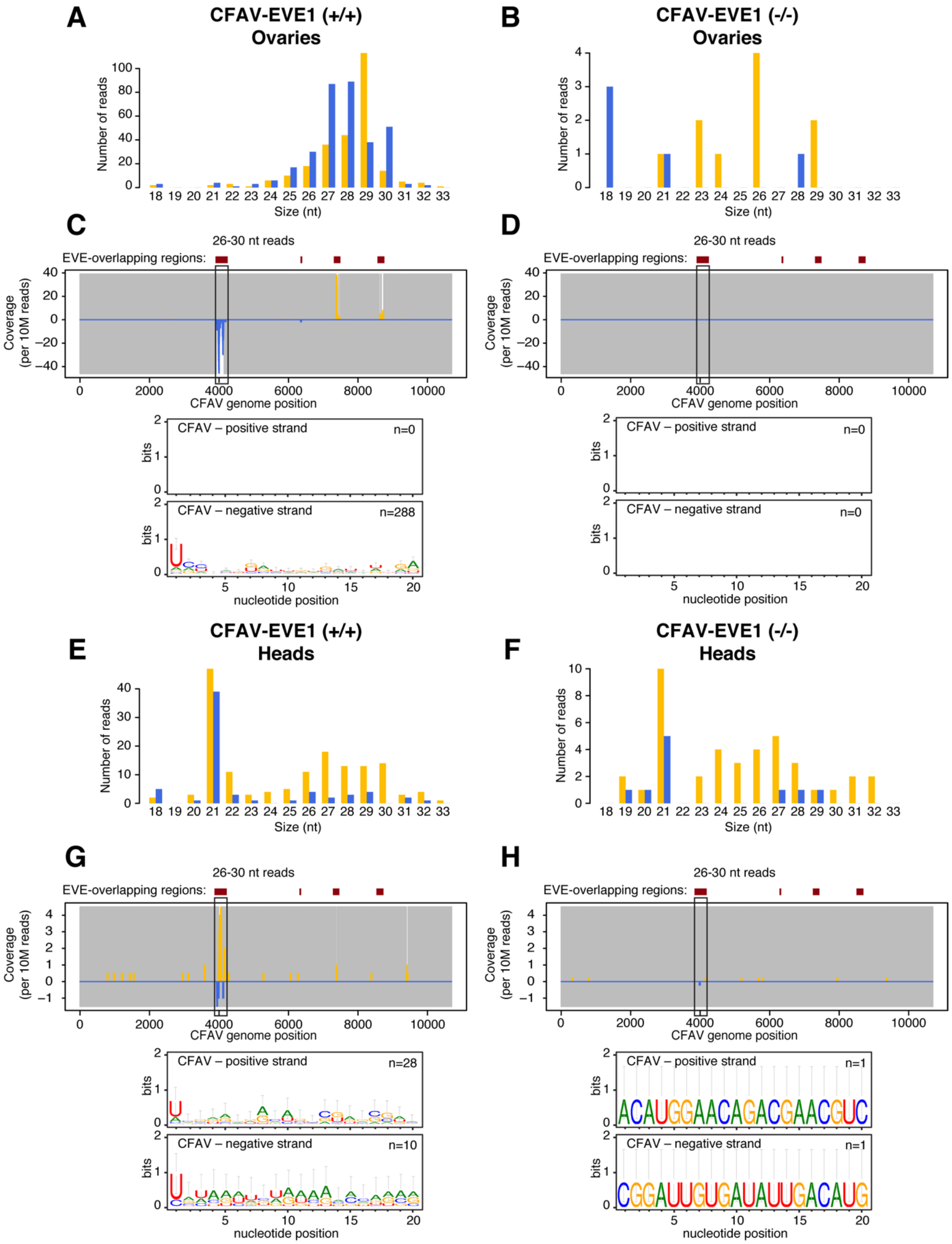


**Figure S4. Small RNA profiles of CFAV-EVE1 (+/+) and (-/-) mock-inoculated *Aedes aegypti* lines. Related to Figure 4.**

Size distribution of sRNAs mapping to the CFAV genome in ovaries (**A-B**) and heads (**E-F**) from experimentally mock-infected CFAV-EVE1 (+/+) (**A,E**) and CFAV-EVE1 (-/-) (**B,F**) mosquitoes 7 days post injection. Analysis of CFAV-derived piRNAs in ovaries (**C-D**) and heads (**G-H**) from experimentally mock-infected CFAV-EVE1 (+/+) (**C,G**) and CFAV-EVE1 (-/-) (**D,H**) mosquitoes 7 days post injection. Mapping (top) and sequence logos (bottom) of 26-30 nt sRNAs. Positive- and negative-sense reads with respect to the reference CFAV genome are shown in yellow and blue, respectively. Uncovered nucleotides are represented by gray lines.

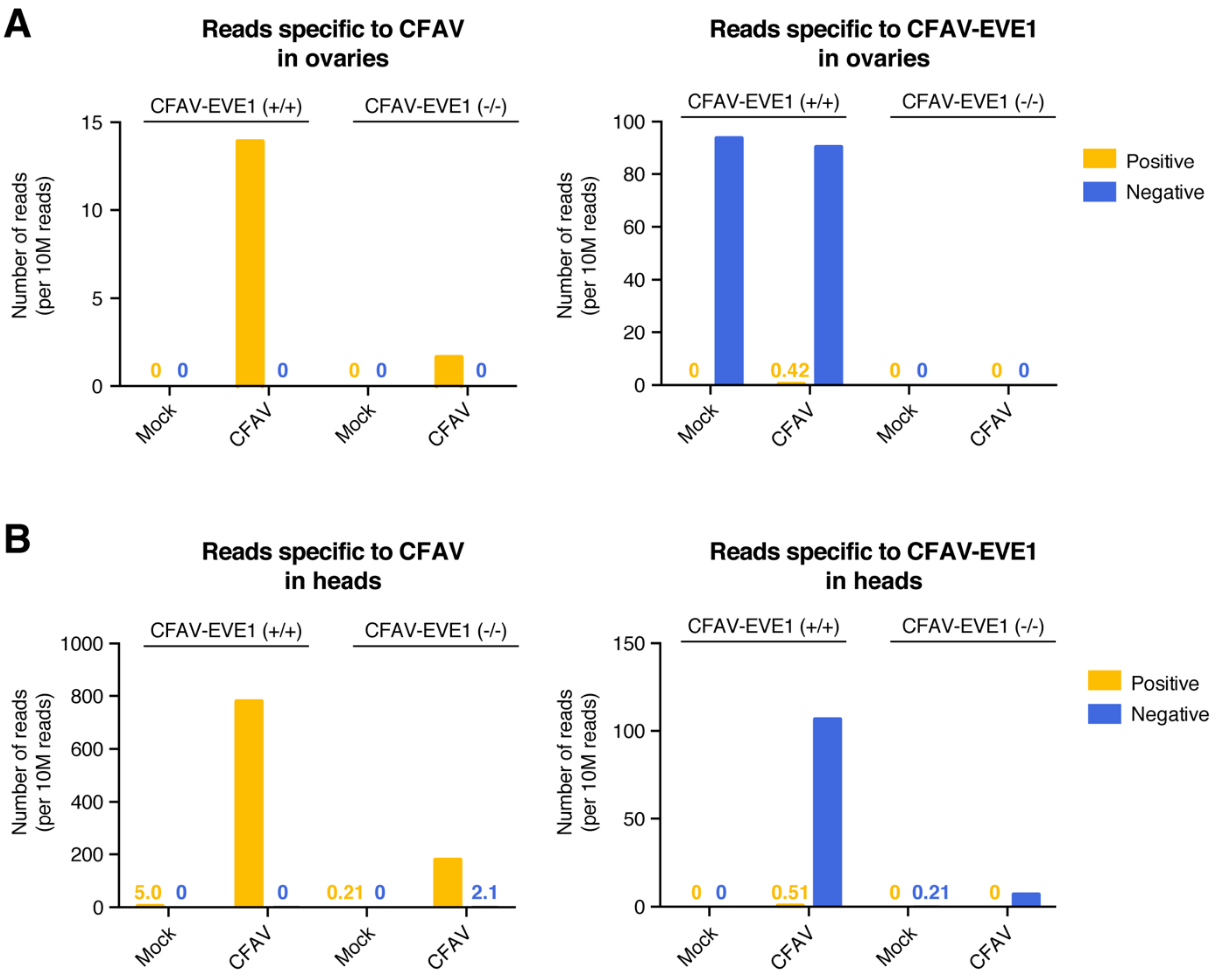


**Figure S5. Target specificity of the piRNAs upon CFAV infection in CFAV-EVE1 (+/+) and (-/-) *Aedes aegypti* lines. Related to Figure 4.**

Number of 26-30 nt sRNA reads unambiguously mapping to the CFAV-KPP genome (left) or to the CFAV-EVE1 locus (right) in ovaries (**A**) and heads (**B**) of mosquitoes from the CFAV-EVE1 (+/+) line and CFAV-EVE1 (-/-) line 7 days after mock-infection or CFAV infection. Yellow and blue colors represent positive-sense and negative-sense reads, respectively. For visual clarity, the normalized number of reads is shown above the bars when the number of reads is <1% of the maximum value.
